# Comparing methods for detecting multilocus adaptation with multivariate genotype-environment associations

**DOI:** 10.1101/129460

**Authors:** Brenna R. Forester, Jesse R. Lasky, Helene H. Wagner, Dean L. Urban

## Abstract

Identifying adaptive loci can provide insight into the mechanisms underlying local adaptation. Genotype-environment association (GEA) methods, which identify these loci based on correlations between genetic and environmental data, are particularly promising. Univariate methods have dominated GEA, despite the high dimensional nature of genotype and environment. Multivariate methods, which analyze many loci simultaneously, may be better suited to these data since they consider how sets of markers covary in response to environment. These methods may also be more effective at detecting adaptive processes that result in weak, multilocus signatures. Here, we evaluate four multivariate methods, and five univariate and differentiation-based approaches, using published simulations of multilocus selection. We found that Random Forest performed poorly for GEA. Univariate GEAs performed better, but had low detection rates for loci under weak selection. Constrained ordinations showed a superior combination of low false positive and high true positive rates across all levels of selection. These results were robust across the demographic histories, sampling designs, sample sizes, and levels of population structure tested. The value of combining detections from different methods was variable, and depended on study goals and knowledge of the drivers of selection. Reanalysis of genomic data from gray wolves highlighted the unique, covarying sets of adaptive loci that could be identified using redundancy analysis, a constrained ordination. Although additional testing is needed, this study indicates that constrained ordinations are an effective means of detecting adaptation, including signatures of weak, multilocus selection, providing a powerful tool for investigating the genetic basis of local adaptation.

## Introduction

Analyzing genomic data for loci underlying local adaptation has become common practice in evolutionary and ecological studies (Hoban *et al.,* 2016). These analyses can help identify mechanisms of local adaptation and inform management decisions for agricultural, natural resources, and conservation applications. Genotype-environment association (GEA) approaches are particularly promising for detecting these loci (Rellstab *et al.* 2015). Unlike differentiation outlier methods, which identify loci with strong allele frequency differences among populations, GEA approaches identify adaptive loci based on associations between genetic data and environmental variables hypothesized to drive selection. Benefits of GEA include the option of using individual-based (as opposed to population-based) sampling and the ability to make explicit links to the ecology of organisms by including relevant predictors. The inclusion of predictors can also improve power and allows for the detection of selective events that do not produce high genetic differentiation among populations (De Mita *et al.,* 2013; de Villemereuil *et al.,* 2014; Rellstab *et al.,* 2015).

Univariate statistical methods have dominated GEA since their first appearance (Mitton *et al.,* 1977). These methods test one locus and one predictor variable at a time, and include generalized linear models (e.g. Joost et al. 2007; Stucki et al. 2016), variations on linear mixed effects models (e.g. Coop et al. 2010; Frichot et al. 2013; Yoder et al. 2014; Lasky et al. 2014), and non-parametric approaches (e.g. partial Mantel, Hancock et al. 2011). While these methods perform well, they can produce elevated false positive rates in the absence of correction for multiple comparisons, an issue of increased importance with large genomic data sets. Corrections such as Bonferroni can be overly conservative (potentially removing true positive detections), while alternative correction methods, such as false discovery rate (FDR, Benjamini & Hochberg 1995), rely on an assumption of a null distribution of *p*-values, which may often be violated for empirical data sets. While these issues should not discourage the use of univariate methods (though corrections should be chosen carefully, see François *et al.* (2016) for a recent overview), other analytical approaches may be better suited to the high dimensionality of modern genomic data sets.

In particular, multivariate approaches, which analyze many loci simultaneously, are well suited to data sets comprising hundreds of individuals sampled at many thousands of genetic markers. Compared to univariate methods, these approaches are thought to more effectively detect multilocus selection since they consider how groups of markers covary in response to environmental predictors (Rellstab et al. 2015). This is important because many adaptive processes are expected to result in weak, multilocus molecular signatures due to selection on standing genetic variation, recent/contemporary selection that has not yet led to allele fixation, and conditional neutrality (Yeaman & Whitlock, 2011; Le Corre & Kremer, 2012; Savolainen *et al.,* 2013; Tiffin & Ross-Ibarra, 2014). Identifying the relevant patterns (e.g., coordinated shifts in allele frequencies across many loci) that underlie these adaptive processes is essential to both improving our understanding of the genetic basis of local adaptation, and advancing applications of these data for management, such as conserving the evolutionary potential of species (Savolainen *et al.,* 2013; Harrisson *et al.,* 2014; Lasky *et al.,* 2015). While multivariate methods may, in principle, be better suited to detecting these shared patterns of response, they have not yet been tested on common data sets simulating multilocus adaptation, limiting confidence in their effectiveness on empirical data.

Here we evaluate a set of these methods, using published simulations of multilocus selection (Lotterhos & Whitlock, 2014, 2015). We compare power using empirical *p*-values, and evaluate false positive rates based on cutoffs used in empirical studies. We follow up with a test of three of these methods on their ability to detect weak multilocus selection, as well as an assessment of the common practice of combining detections across multiple tests. We investigate the effects of correction for population structure in one ordination method, and follow up with an application of this test to an empirical data set from gray wolves. We find that the constrained ordinations we tested maintain the best balance of true and false positive rates across a range of demographies, sampling designs, sample sizes, and selection levels, and can provide unique insight into the processes driving selection and the multilocus architecture of local adaptation.

## Methods

### Multivariate approaches to GEA

Multivariate statistical techniques, including ordinations such as principal components analysis (PCA), have been used to analyze genetic data for over fifty years (Cavalli-Sforza, 1966).Indirect ordinations like PCA (which do not use predictors) use patterns of association within genetic data to find orthogonal axes that fully decompose the genetic variance. Constrained ordinations extend this analysis by restricting these axes to combinations of supplied predictors (Jombart *et al.,* 2009; Legendre & Legendre, 2012). When used as a GEA, a constrained ordination is essentially finding orthogonal sets of loci that covary with orthogonal multivariate environmental patterns. By contrast, a univariate GEA is testing for single locus relationships with single environmental predictors. The use of constrained ordinations in GEA goes back as far as Mulley et al. (1979), with more recent applications to genomic data sets in Lasky et al. (2012), Forester et al. (2016), and Brauer et al. (2016). In this analysis, we test two promising constrained ordinations, redundancy analysis (RDA) and distance-based redundancy analysis (dbRDA). We also test an extension of RDA that uses a preliminary step of summarizing the genetic data into sets of covarying markers (Bourret *et al.,* 2014). We do not include canonical correspondence analysis, a constrained ordination that is best suited to modeling unimodal responses, although this method has been used to analyze microsatellite data sets (e.g. Angers et al. 1999; Grivet et al. 2008).

Random Forest (RF) is a machine learning algorithm that is designed to identify structure in complex data and generate accurate predictive models. It is based on classification and regression trees (CART), which recursively partition data into response groups based on splits in predictors variables. CART models can capture interactions, contingencies, and nonlinear relationships among variables, differentiating them from linear models (De’ath & Fabricius, 2000). RF reduces some of the problems associated with CART models (e.g. overfitting and instability) by building a “forest” of classification or regression trees with two layers of stochasticity: random bootstrap sampling of the data, and random subsetting of predictors at each node (Breiman, 2001). This provides a built-in assessment of predictive accuracy (based on data left out of the bootstrap sample) and variable importance (based on the change in accuracy when covariates are permuted). For GEA, variable importance is the focal statistic, where the predictor variables used at each split in the tree are molecular markers, and the goal is to sort individuals into groups based on an environmental category (classification) or to predict home environmental conditions (regression). Markers with high variable importance are best able to sort individuals or predict environments. RF has been used in a number of recent GEA and GWAS studies (e.g. Holliday et al. 2012; Brieuc et al. 2015; Pavey et al. 2015; Laporte et al. 2016), but has not yet been tested in a GEA simulation framework.

We compare these multivariate methods to the two differentiation-based and three univariate GEA methods tested by Lotterhos & Whitlock (2015): the X^T^X statistic from Bayenv2 (Günther & Coop, 2013), PCAdapt (Duforet-Frebourg *et al.,* 2014), latent factor mixed models (LFMM, Frichot *et al.* 2013), and two GEA-based statistics (Bayes factors and Spearman’s ρ) from Bayenv2. We also include generalized linear models (GLM), a regression-based GEA that does not use a correction for population structure.

### GEA implementation

#### Constrained ordinations

We tested RDA and dbRDA as implemented by Forester et al. (2016). RDA is a two-step process in which genetic and environmental data are analyzed using multivariate linear regression, producing a matrix of fitted values. Then PCA of the fitted values is used to produce canonical axes, which are linear combinations of the predictors. We centered and scaled genotypes for RDA (i.e., mean = 0, s = 1; see Jombart *et al.* 2009 for a discussion of scaling genetic data for ordinations). Distance-based redundancy analysis is similar to RDA but allows for the use of non-Euclidian dissimilarity indices. Whereas RDA can be loosely considered as a PCA constrained by predictors, dbRDA is analogous to a constrained principal coordinate analysis (PCoA, or a PCA on a non-Euclidean dissimilarity matrix). For dbRDA, we calculated the distance matrix using Bray-Curtis dissimilarity (Bray & Curtis, 1957), which quantifies the dissimilarity among individuals based on their multilocus genotypes (equivalent to one minus the proportion of shared alleles between individuals). For both methods, SNPs are modeled as a function of predictor variables, producing as many constrained axes as predictors. We identified outlier loci on the constrained ordination axes based on the “locus score”, which represent the coordinates/loading of each locus in the ordination space. We use *rda* for RDA and *capscale* for dbRDA in the vegan, v. 2.3-5 package (Oksanen *et al.,* 2013) in R v. 3.2.5 (R Development Core Team, 2015) for this and all subsequent analyses.

#### Redundancy analysis of components

This method, described by Bourret *et al.* (2014), differs from the approaches described above in using a preliminary step that summarizes the genotypes into sets of covarying markers, which are then used as the response in RDA. The idea is to identify from these sets of covarying loci only the groups that are most strongly correlated with environmental predictors. We began by ordinating SNPs into principal components (PCs) using *prcomp* in R on the scaled data, producing as many axes as individuals. Following Bourret et al. (2014), we used parallel analysis (Horn, 1965) to determine how many PCs to retain. Parallel analysis is a Monte Carlo approach in which the eigenvalues of the observed components are compared to eigenvalues from simulated data sets that have the same size as the original data. We used 1,000 random data sets to generate the distribution under the null hypothesis and retained components with eigenvalues greater than the 99^th^ percentile of the eigenvalues of the simulated data (i.e., a significance level of 0.01), using the hornpa package, v. 1.0 (Huang, 2015).

Next, we applied a varimax rotation to the PC axes, which maximizes the correlation between the axes and the original variables (in this case, the SNPs). Note that once a rotation is applied to the PC axes, they are no longer “principal” components (i.e. axes associated with an eigenvalue/variance), but simply components. We then used the retained components as dependent variables in RDA, with environmental variables used as predictors. Next, components that were significantly correlated with the constrained axis were retained. Significance was based on a cutoff (alpha = 0.05) corrected for sample sizes using a Fisher transformation as in Bourret et al. (2014). Finally, SNPs were correlated with these retained components to determine outliers. We call this approach redundancy analysis of components (cRDA).

#### Random Forest

The Random Forest approach implemented here builds off of work by Goldstein et al. (2010), Holliday et al. (2012), and Brieuc et al. (2015). This three-step approach is implemented separately for each predictor variable. The environmental variable used in this study was continuous, so RF models were built as regression trees. For categorical predictors (e.g. soil type) classification trees would be used, which require a different parameterization (important recommendations for this case are provided in Goldstein et al. 2010).

First, we tuned the two main RF parameters, the number of trees *(ntrees)* and the number of predictors sampled per node (*mtry*). We tested a range of values for *ntrees* in a subset of the simulations, and found that 10,000 trees were sufficient to stabilize variable importance (note that variable importance requires a larger number of trees for convergence than error rates, Goldstein et al. 2010). We used the default value of *mtry* for regression (number of predictors/3, equivalent to ∼3,330 SNPs in this case) after checking that increasing *mtry* did not substantially change variable importance or the percent variance explained. In a GEA/GWAS context, larger values of *mtry* reduce error rates, improve variable importance estimates, and lead to greater model stability (Goldstein *et al.* 2010).

Because RF is a stochastic algorithm, it is best to use multiple runs, particularly when variable importance is the parameter of interest (Goldstein *et al.,* 2010). We begin by building three full RF models using all SNPs as predictors, saving variable importance as mean decrease in accuracy for each model. Next, we sampled variable importance from each run with a range of cutoffs, pulling the most important 0.5%, 1.0%, 1.5%, and 2.0% of loci. These values correspond to approximately 50/100/150/200 loci that have the highest variable importance. For each cutoff, we then created three additional RF models, using the average percent variance explained across runs to determine the best starting number of important loci for step 3. This step removes clearly unimportant loci from further consideration (i.e. “sparsity pruning”, Goldstein et al. 2010).

Third, we doubled the best starting number of loci from step 2; this is meant to accommodate loci that may have low marginal effects (Goldstein et al. 2010). We then built three RF models with these loci, and recorded the mean variance explained. We removed the least important locus in each model, and recalculated the RF models and mean variance explained. This procedure continues until two loci remain. The set of loci that explain the most variance are the final candidates. Candidates are then combined across runs to identify outliers.

#### Differentiation-based and univariate GEA methods

For the two differentiation-based and the Bayenv2-based GEA methods, we compared power directly from the results provided in Lotterhos & Whitlock (2015). PCAdapt is a differentiation-based method that concurrently identifies outlier loci and population structure using latent factors (Duforet-Frebourg *et al.,* 2014). The X^T^X statistic from Bayenv2 (Günther & Coop, 2013) is an F_ST_ analog that uses a covariance matrix to control for population structure. The two Bayenv2 GEA statistics (Bayes factors and Spearman’s ρ) also use the covariance matrix to control for population structure, while identifying candidate loci based on log-transformed Bayes factors and nonparametric correlations, respectively. Details on these methods and their implementation are provided in Lotterhos & Whitlock (2015).

We reran latent factor mixed models, a GEA approach that controls for population structure using latent factors, using updated parameters as recommended by the authors (O. François, pers. comm.). We tested values of *K* (the number of latent factors) ranging from one to 25 using a sparse nonnegative matrix factorization algorithm (Frichot *et al.,* 2014), implemented as function *snmf* in the package LEA, v. 1.2.0 (Frichot & François, 2015). We plotted the cross-entropy values and selected *K* based on the inflection point in these plots; when the inflection point was not clear, we used the value where additional cross-entropy loss was minimal. We parameterized LFMM models with this best estimate of *K*, and ran each model ten times with 5,000 iterations and a burnin of 2,500. We used the median of the squared z-scores to rank loci and calculate a genomic inflation factor (GIF) to assess model fit (Frichot & François, 2015; François *et al.,* 2016). The GIF is used to correct for inflation of z-scores at each locus, which can occur when population structure or other confounding factors are not sufficiently accounted for in the model (François *et al.* 2016). The GIF is calculated by dividing the median of the squared z-scores by the median of the chi-squared distribution. We used the LEA and qvalue, v. 2.2.2 (Storey *et al.,* 2015) packages in R. Full K and GIF results are presented in Table S1. Finally, we ran generalized linear models (GLM) on individual allele counts using a binomial family and logistic link function for comparison with LFMM; GIF results are presented in Table S1.

### Simulations

We used a subset of simulations published by Lotterhos & Whitlock (2014, 2015). Briefly, four demographic histories are represented in these data, each with three replicated environmental surfaces (Fig. S1): an equilibrium island model (IM), equilibrium isolation by distance (IBD), and nonequilibrium isolation by distance with expansion from one (1R) or two (2R) refugia. In all cases, demography was independent of selection strength, which is analogous to simulating soft selection (Lotterhos & Whitlock, 2014). Haploid, biallelic SNPs were simulated independently, with 9,900 neutral loci and 100 under selection. Note that haploid SNPs will yield half the information content of diploid SNPs (Lotterhos & Whitlock 2015). The mean of the environmental/habitat parameter had a selection coefficient equal to zero and represented the background across which selective habitat was patchily distributed (Fig. S1). Selection coefficients represent a proportional increase in fitness of alleles in response to habitat, where selection is increasingly positive as the environmental value increases from the mean, and increasingly negative as the value decreases from the mean (Lotterhos & Whitlock 2014, Fig.S1). This landscape emulates a weak cline, with a north-south trend in the selection surface. Of the 100 adaptive loci, most were under weak selection. For the IBD scenarios, selection coefficients were 0.001 for 40 loci, 0.005 for 30 loci, 0.01 for 20 loci, and 0.1 for 10 loci. For the 1R, 2R, and IM scenario, selection coefficients were 0.005 for 50 loci, 0.01 for 33 loci, and 0.1 for 17 loci. Note that realized selection varied across demographies, so results across demographic histories are not directly comparable (Lotterhos & Whitlock 2015).

We used the following sampling strategies and sample sizes from Lotterhos & Whitlock (2015): random, paired, and transect strategies, with 90 demes sampled, and 6 or 20 individuals sampled per deme. Paired samples (45 pairs) were designed to maximize environmental differences between locations while minimizing geographic distance; transects (nine transects with ten locations) were designed to maximize environmental differences at transect ends (Lotterhos & Whitlock 2015). Overall, we used 72 simulations for testing. We assessed trend in neutral loci using linear models of allele frequencies within demes as a function of coordinates. We evaluated the strength of local adaptation using linear models of allele frequencies within demes as a function of environment. Note that the Lotterhos & Whitlock (2014, 2015) simulations assigned SNP genotypes to individuals within a population sequentially (i.e., the first few individuals would all get the same allele until its target frequency was reached, the remaining individuals would get the other allele). This creates artifacts (e.g., artificially low observed heterozygosity) and may affect statistical error rates when subsampling individuals or performing analyses at the individual level. As recommended by K. Lotterhos (pers. comm.), we avoided these problems by randomizing allele counts for each SNP among individuals within each population. The habitat surface, which imposed a continuous selective gradient on nonneutral loci, was used as the environmental predictor.

### Evaluation statistics

In order to equitably compare power (true positive detections out of the number of loci under selection) across these methods, we calculated empirical *p*-values using the method of Lotterhos & Whitlock (2015). In this approach, we first built a null distribution based on the test statistics of all neutral loci, and then generated a *p*-value for each selected locus based on its cumulative frequency in the null distribution. We then converted empirical *p*-values to *q*-values to assess significance, using the same *q*-value cutoff (0.01) as Lotterhos & Whitlock (2015). We used code provided by K. Lotterhos to calculate empirical *p*-values (code provided in Supplemental Information).

Because false positive rates (FPRs) are not very informative for empirical *p*-values (rates are universally low, see Lotterhos & Whitlock 2015 for a discussion), we applied cutoffs (e.g. thresholds for statistical significance) to assess both true and false positive rates across methods. While power is important, determining FPRs is also an essential component of assessing method performance, since high power achieved at the cost of high FPRs is problematic. Because cutoffs differ across methods, we tested a range of commonly used thresholds for each method and chose the approach that performed the best (i.e., best balance of TPR and FPR). Note that cutoffs can be adjusted for empirical studies based on research goals and the tolerance for TP and FP detections. For each cutoff tested, we calculated the TPR as the number of correct positive detections out of the number possible, and the FPR as the number of incorrect positive detections out of 9900 possible. For the main text, we present results from the best cutoff for each method; full results for all cutoffs tested are presented in the Supplemental Information. For constrained ordinations (RDA and dbRDA) we identified outliers as SNPs with a locus score +/− 2.5 and 3 SD from the mean score of each constrained axis. For cRDA, we used cutoffs for SNP-component correlations of alpha = 0.05, 0.01, and 0.001, corrected for sample sizes using a Fisher transformation as in Bourret et al. (2014). For GLM and LFMM, we compared two Bonferroni-corrected cutoffs (0.05 and 0.01) and a FDR cutoff of 0.1.

### Weak selection

We compared the best-performing multivariate methods (RDA, dbRDA, and cRDA) for their ability to detect signals of weak selection (s = 0.005 and s = 0.001). All tests were performed as described above, after removing loci under strong (s = 0.1) and moderate (s = 0.01) selection from the simulation data sets. The number of loci under selection in these cases ranged from 43 to 76.

### Combining detections

We compared the effects of combining detections (i.e., looking for overlap) using cutoff results from two of the best-performing methods, RDA and LFMM. We also included a scenario in which a second, uninformative predictor (the x-coordinate of each individual) is included in the RDA and LFMM tests. This predictor is analogous to including an environmental variable hypothesized to drive selection that covaries with longitude.

### Correction for population structure in RDA

To determine how explicit modeling of population structure affects the performance of the best-performing multivariate method, RDA, we accounted for population structure using three approaches: (1) partialling out significant spatial eigenvectors not correlated with the habitat predictor, (2) partialling out all significant spatial eigenvectors, and (3) partialling out ancestry coefficients. The spatial eigenvector procedure uses Moran eigenvector maps (MEM) as spatial predictors in a partial RDA. MEMs provide a decomposition of the spatial relationships among sampled locations based on a spatial weighting matrix (Dray *et al.,* 2006). We used spatial filtering to determine which MEMs to include in the partial analyses (Dray *et al.,* 2012). Briefly, this procedure begins by applying a principal coordinate analysis (PCoA) to the genetic distance matrix, which we calculated using Bray-Curtis dissimilarity. We used the broken-stick criterion (Legendre & Legendre, 2012) to determine how many genetic PCoA axes to retain. Retained axes were used as the response in a full RDA, where the predictors included all MEMs. Forward selection (Blanchet *et al.,* 2008) was used to reduce the number of MEMs, using the full RDA adjusted R^2^ statistic as the threshold. In the first approach, retained MEMs that were significantly correlated with environmental predictors were removed (alpha = 0.05/number of MEMs), and the remaining set of significant MEMs were used as conditioning variables in RDA. Note that this approach will be liberal in removing MEMs correlated with environment. In the second approach, all significant MEMs were used as conditioning variables, the most conservative use of MEMs. We used the spdep, v. 0.6-9 (Bivand *et al.,* 2013) and adespatial, v. 0.0-7 (Dray *et al.,* 2016) packages to calculate MEMs. For the third approach, we used individual ancestry coefficients as conditioning variables. We used function *snmf* in the LEA package to estimate individual ancestry coefficients, running five replicates using the best estimate of K, and extracting individual ancestry coefficients from the replicate with the lowest cross-entropy.

### Empirical data set

To provide an example of the use and interpretation of RDA as a GEA, we reanalyzed data from 94 North American gray wolves (*Canis lupus*) sampled across Canada and Alaska at 42,587 SNPs (Schweizer *et al.,* 2016). These data show similar global population structure to the simulations analyzed here: wolf data Fst = 0.09; average simulation Fst = 0.05. We reduced the number of environmental covariates originally used by Schweizer *et al.* (2016) from 12 to eight to minimize collinearity among them (e.g., |r| < 0.7). One predictor, land cover, was removed because the distribution of cover types was heavily skewed toward two of the ten types. Missing data levels were low (3.06%). Because RDA requires complete data frames, we imputed missing values by replacing them with the most common genotype across individuals. We identified candidate adaptive loci as SNPs loading +/− 3 SD from the mean loading of significant RDA axes (significance determined by permutation, *p* < 0.05). We then identified the covariate most strongly correlated with each candidate SNP (i.e., highest correlation coefficient), to group candidates by potential driving environmental variables. Annotated code for this example is provided in the Supplementary Information.

## Results

### Empirical *p*-value results

Power across the three ordination techniques was comparable, while power for RF was relatively low (Fig. 1). Ordinations performed best in IBD, 1R, and 2R demographies, with the larger sample size improving power for the IM demography. Within ordination techniques, RDA and cRDA had slightly higher detection rates compared to dbRDA; subsequent comparisons are made using RDA results.

**Figure 1.**
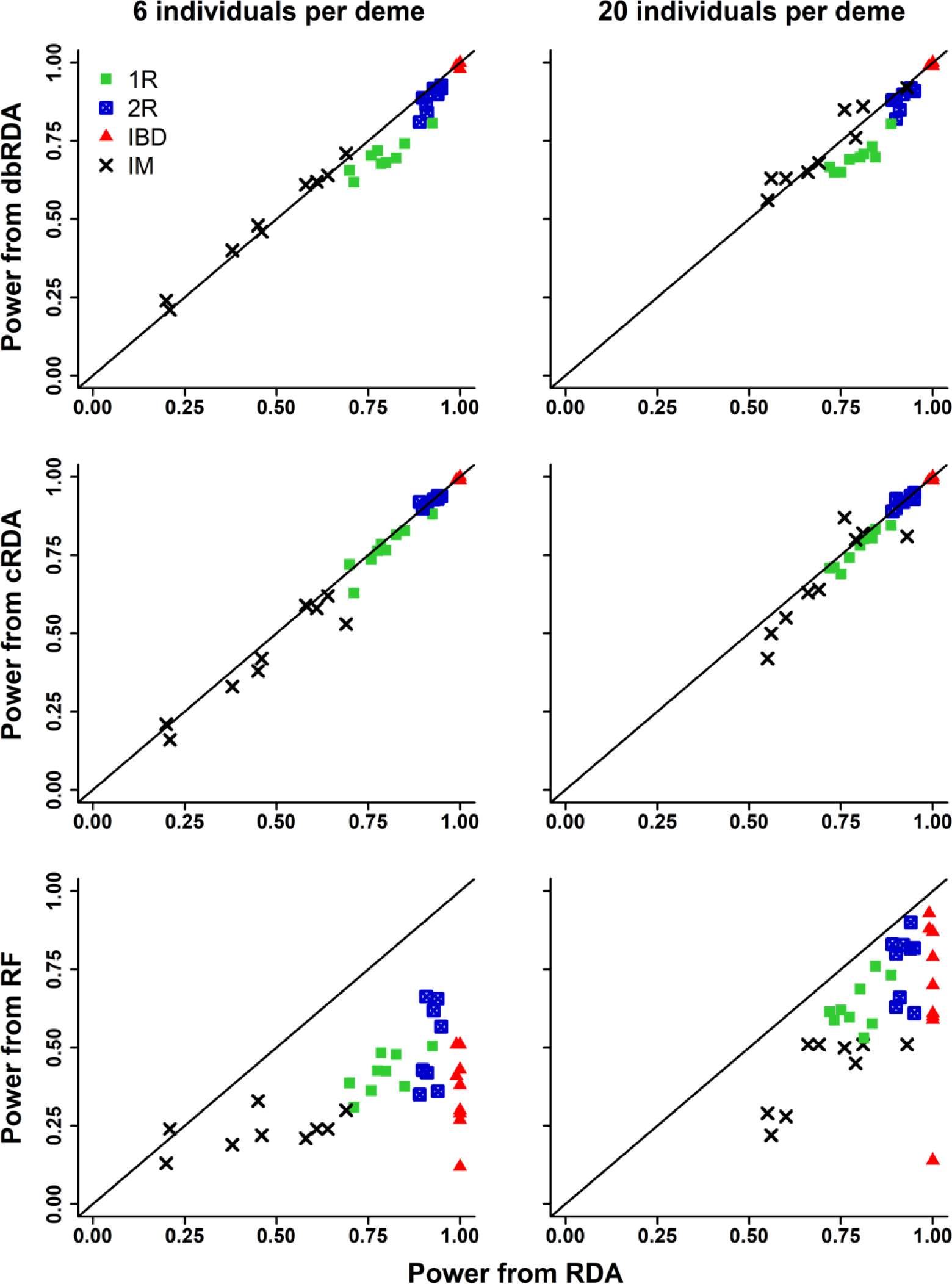
Comparison of power (from empirical *p*-values) from RDA (x-axis) and three other multivariate GEAs (y-axes, rows) for two sample sizes (columns). Points reflect demographies: 1R and 2R = refugial expansion, IBD = equilibrium isolation by distance, IM = equilibrium island model. Some variation within demographies comes from sampling design.

Except for a few cases in the IM demography, the power of RDA was generally higher than univariate GEAs (Fig. 2). Of the univariate methods, GLM had the highest overall power, while LFMM had reduced power for the IBD demography. Power from the Bayes Factor (Bayenv2) was generally lower than RDA across all demographies. Finally, RDA had overall higher power than the two differentiation-based methods (Fig. 3), with the exception of the IBD demography, where power was high for all methods.

**Figure 2.**
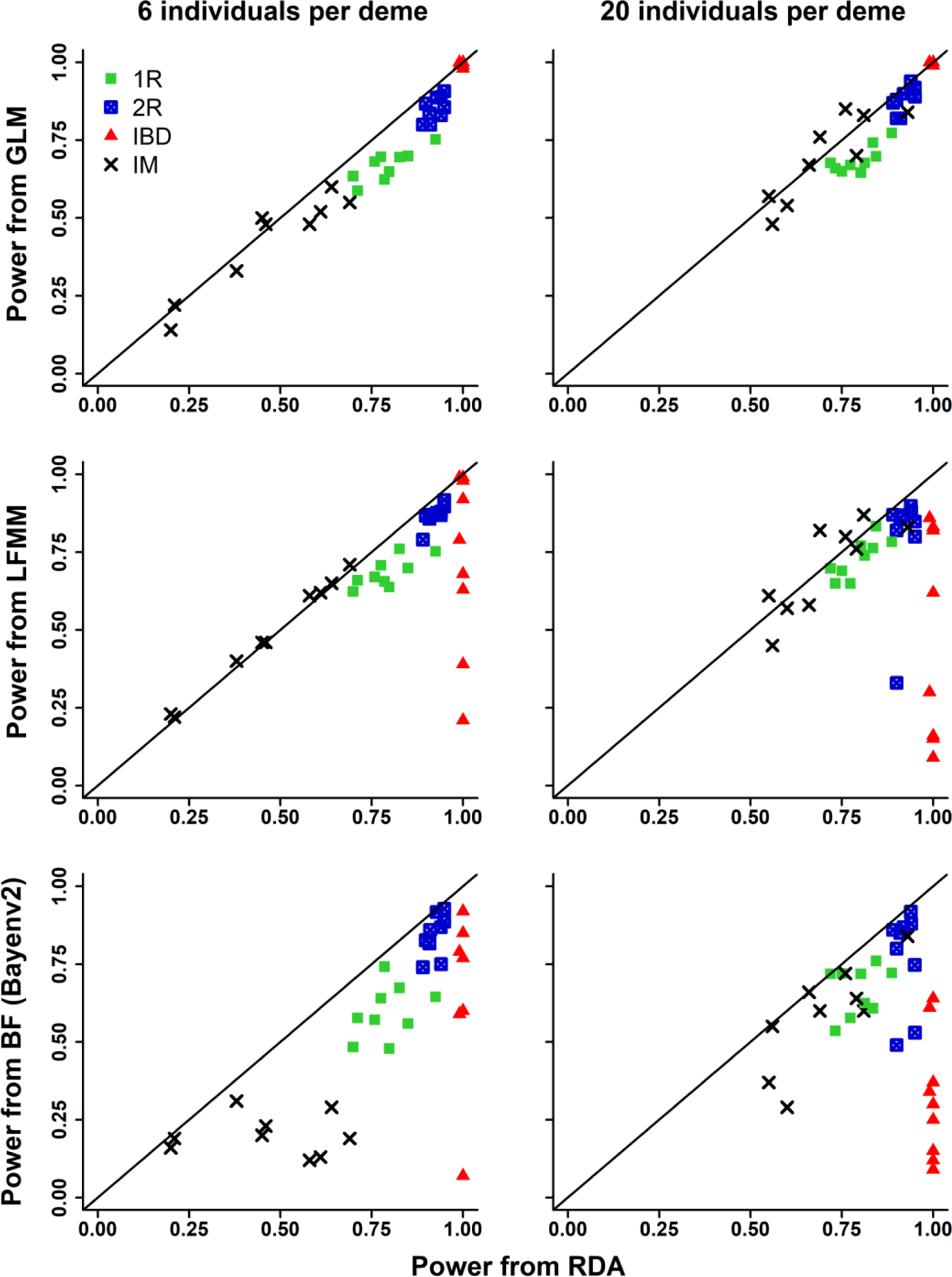
Comparison of power (from empirical *p*-values) from RDA (x-axis) and three univariate GEAs (y-axes, rows) for two sample sizes (columns). Points reflect demographies: 1R and 2R = refugial expansion, IBD = equilibrium isolation by distance, IM = equilibrium island model. Some variation within demographies comes from sampling design.

**Figure 3.**
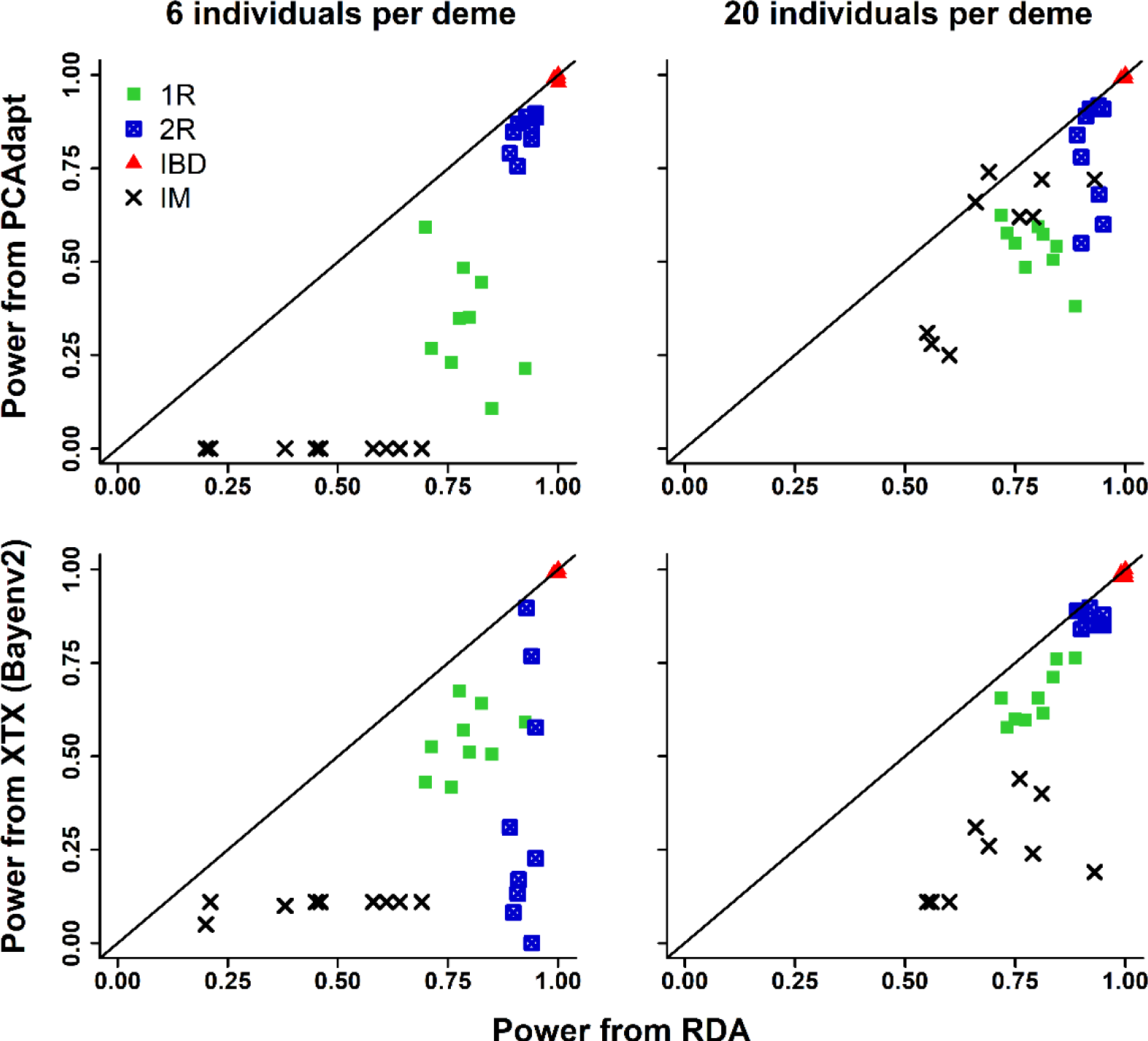
Comparison of power (from empirical *p*-values) from RDA (x-axis) and two differentiation-based outlier detection methods (y-axes, rows) for two sample sizes (columns). Points reflect demographies: 1R and 2R = refugial expansion, IBD = equilibrium isolation by distance, IM = equilibrium island model. Some variation within demographies comes from sampling design.

Among the methods with the highest overall power, all performed well at detecting loci under strong selection (Fig. 4 and S2). Detection rates for loci under moderate and weak selection were highest for ordination methods, with RDA and cRDA having the overall highest detection rates. Detection of moderate and weakly selected loci was lower and more variable for univariate methods, especially LFMM, where detection was dependent on demography and sampling scheme.

**Figure 4.**
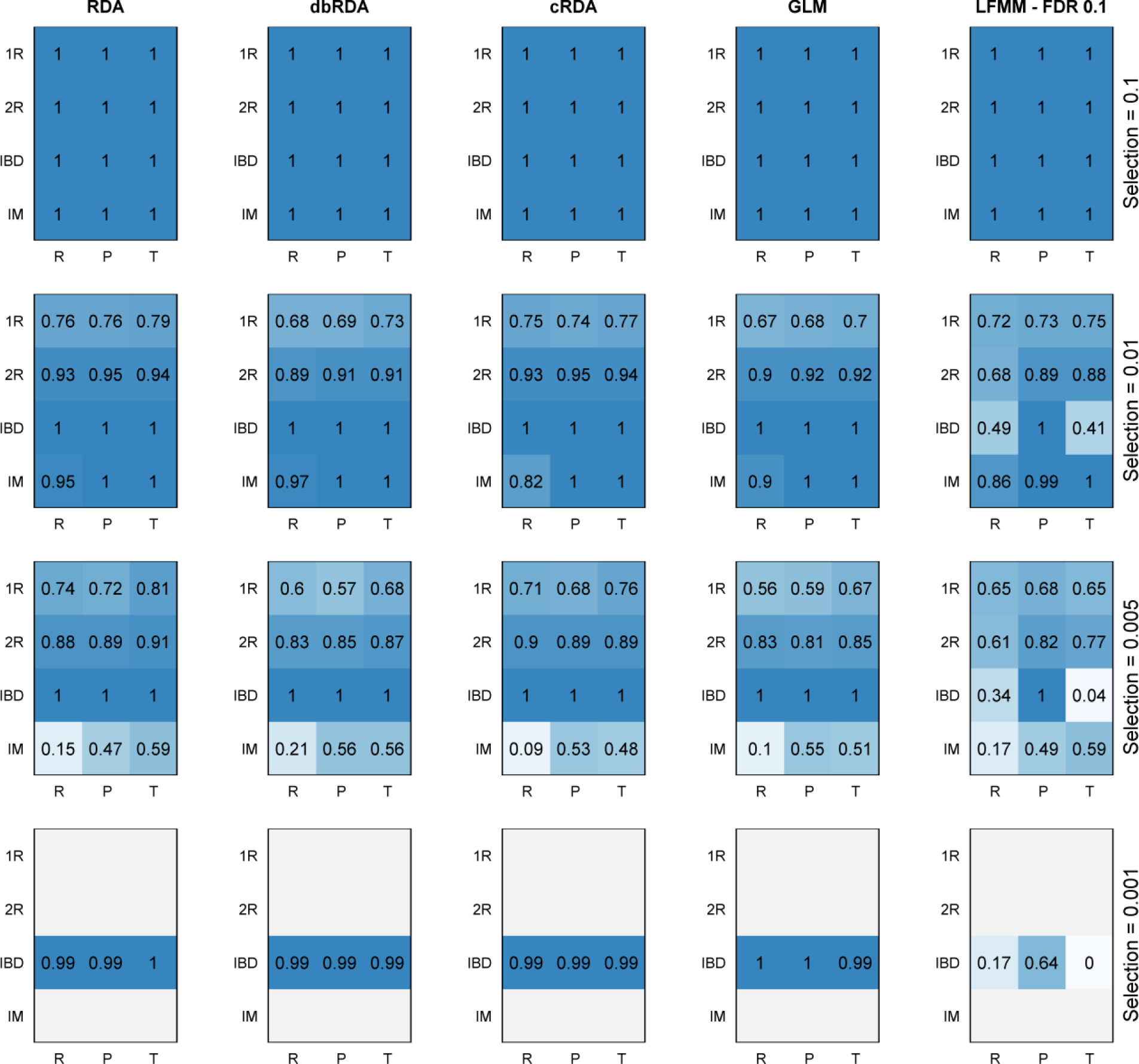
Average power (from empirical *p*-values) for different levels of selection (rows) from five methods (columns) using a sample size of 20 individuals per deme. Each method shows results for different sampling strategies (R = random, P = pairs, T = transects) and demographies (1R and 2R = refugial expansion, IBD = equilibrium isolation by distance, IM = equilibrium island model). Only the IBD demography included very weak selection (s=0.001).

### Weak selection

We compared the three ordination methods for their power to detect only weak loci in the simulations (Fig. 5). Power from RDA was higher when all selected loci were included, especially for the IM demography. Power using only weakly selected loci was comparable between RDA and dbRDA, with power slightly higher for RDA in most cases. cRDA was comparable to RDA for the IBD and 2R demographies, but had very low to no power in the IM demography, and the 1R demography with the larger sample size.

**Figure 5.**
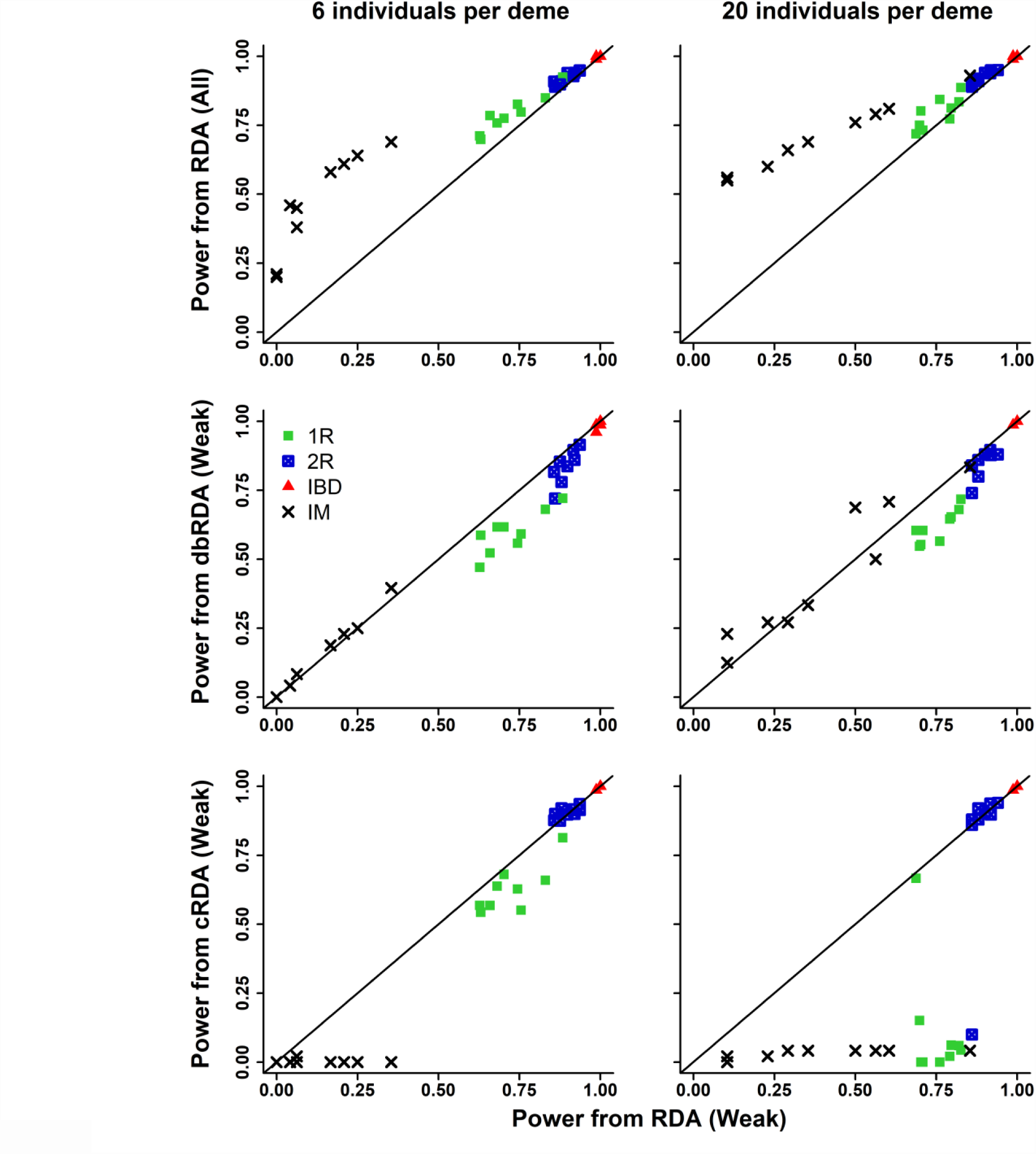
Comparison of power (from empirical *p*-values) from RDA tested on weak selection only (x-axis) and RDA tested on all loci under selection (first row), as well as dbRDA and cRDA tested on weak selection only (second and third rows) for two sample sizes (columns). Points reflect demographies: 1R and 2R = refugial expansion, IBD = equilibrium isolation by distance, IM = equilibrium island model. Some variation within demographies comes from sampling design.

### Cutoff results

We compared cutoff results for the methods with the highest overall power: RDA, dbRDA, cRDA, GLM, and LFMM. The best performing cutoffs were: RDA/dbRDA, +/- 3 SD; cRDA, alpha = 0.001; GLM, Bonferroni = 0.05, and LFMM, FDR = 0.1. We did not choose the FDR cutoff for GLMs since GIFs indicated that the test *p*-values were not appropriately calibrated (i.e., GIFs > 1, Table S1). For some scenarios, LFMM GIFs were less than one (indicating a conservative correction for population structure, Table S1). We reran LFMM models with the best estimate of K minus one (i.e., K-1) to determine if a less conservative correction would influence LFMM results. Because there was no consistent improvement in power or TPR/FPRs using K-1 (Tables S2-S3), all subsequent results refer to LFMM runs using the best estimate of K.

Full cutoff results for each method are presented in the Supplementary Information (Fig. S3-S6). Cutoff FPRs were highest for cRDA and GLM (Fig. 6). By contrast, RDA and dbRDA had mostly zero FPRs, with slightly higher FPRs for LFMM. Within these three low-FPR methods, RDA maintained the highest TPRs, except in the IM demography, where LFMM maintained higher power. LFMM was more sensitive to sampling design than the other methods, with more variation in TPRs across designs.

**Figure 6.**
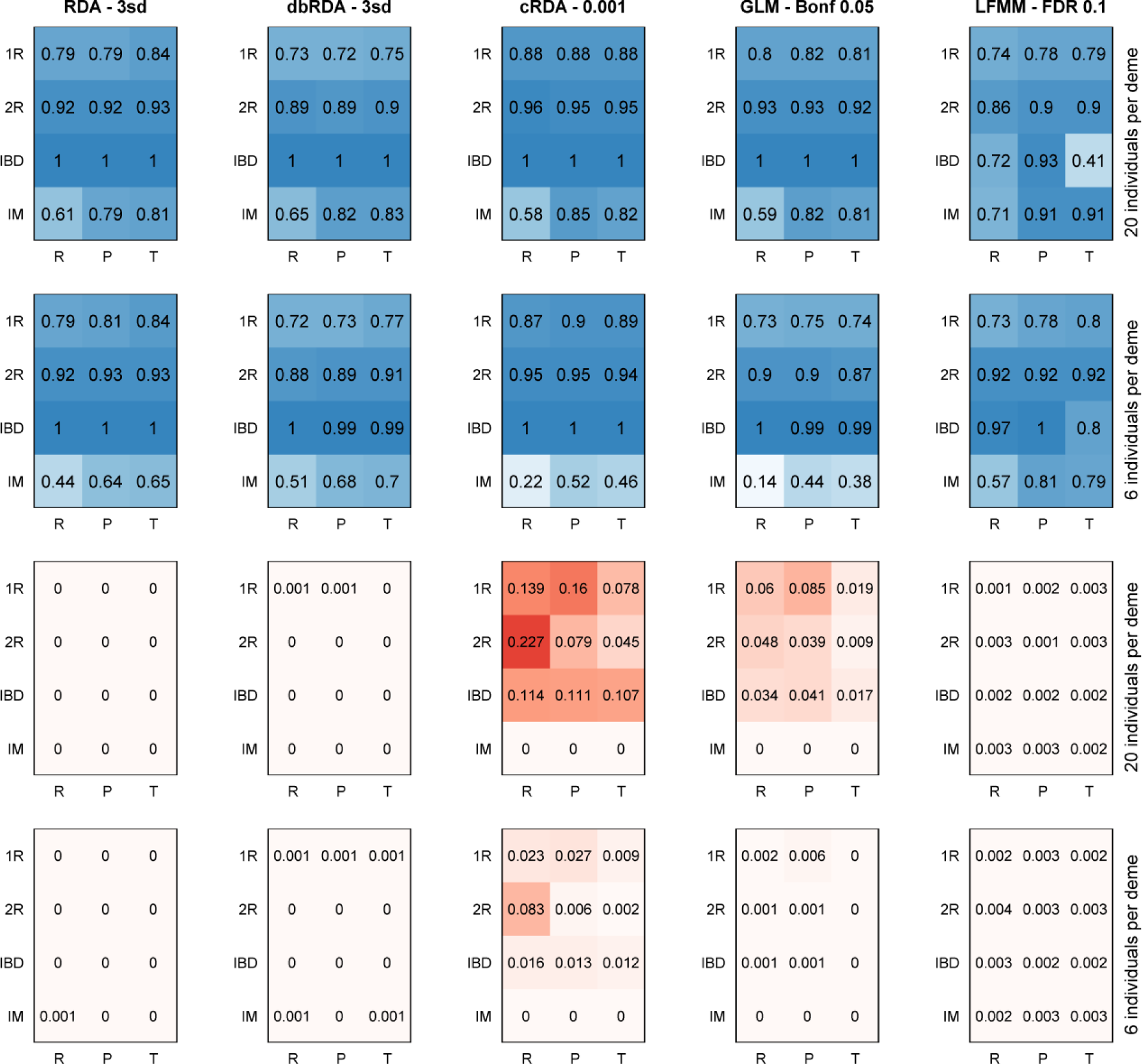
Average true positive (top two rows, in blue) and false positive (bottom two rows, in red) rates from five methods (columns) using the best cutoff for each method. Each method shows results for different sampling strategies (R = random, P = pairs, T = transects), demographies (1R and 2R = refugial expansion, IBD = equilibrium isolation by distance, IM = equilibrium island model), and sample sizes (rows).

### Combining detections

We compared the univariate LFMM and multivariate RDA cutoff results for overlap and differences in their detections using both the habitat predictor only, and the habitat and (uninformative) x-coordinate predictor (Figs. 7 and S7). When the driving environmental predictor is known, RDA detections alone are the best choice, since FPRs are very low and RDA detects a large number of selected loci that are not identified by LFMM (except in the IM demography, Fig. 7a). However, when a noninformative environmental predictor is included, combining test results yields greater overall benefits, since the tests show substantial commonality in TP detections, but show very low commonality in FP detections (Fig. 7b). By retaining only overlapping loci, FPRs are substantially reduced at some loss of power due to discarded RDA (and LFMM in the IM demography) detections.

**Figure 7.**
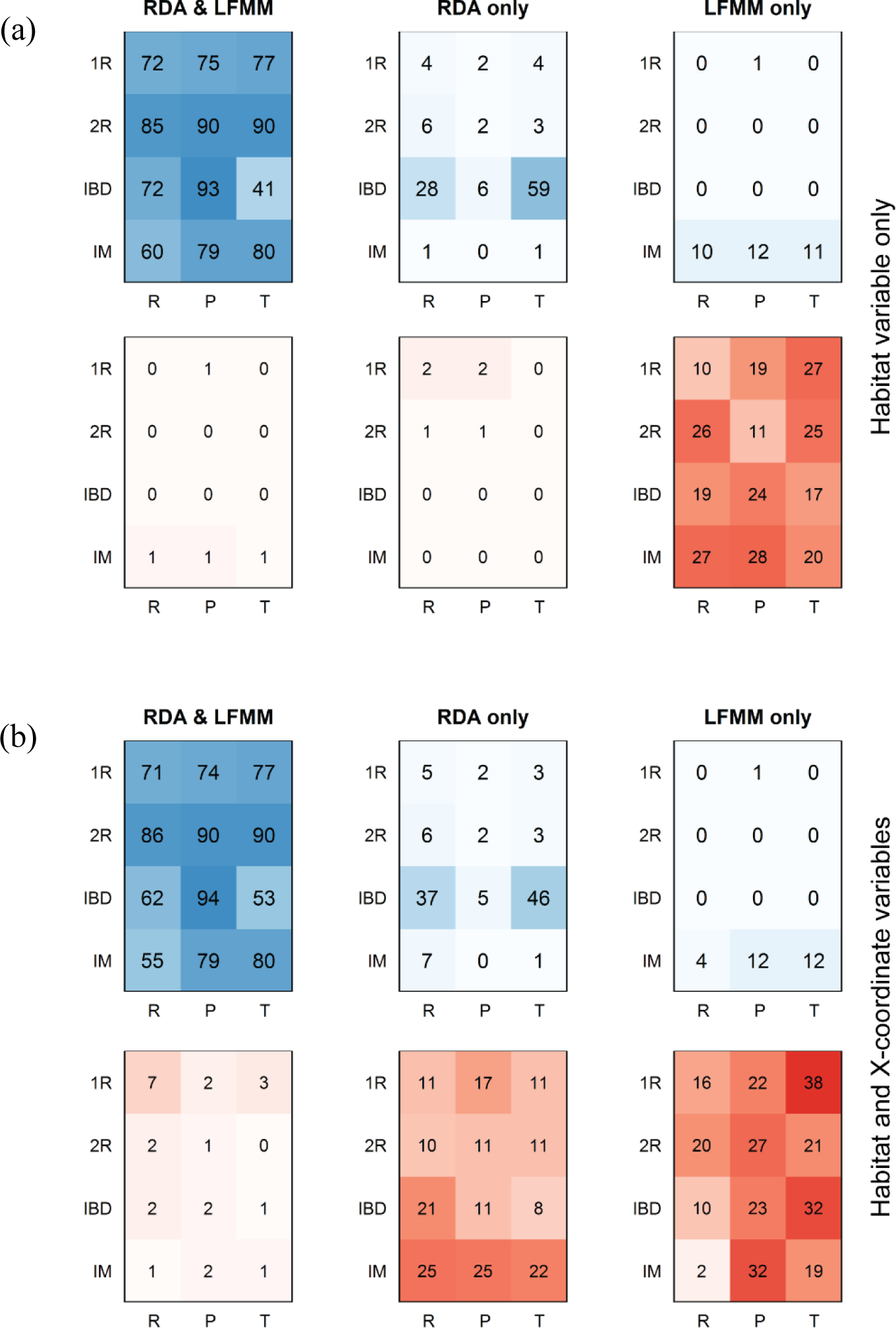
Average counts of true positive (top rows of a and b, in blue) and false positive (bottom rows of a and b, in red) detections for two methods, RDA and LFMM, using their best cutoffs and a sample size of 20 individuals per deme. The first column shows the average number of loci detected by both methods. The second and third columns show the average number of detections that are unique to RDA and LFMM, respectively. (a) Results for GEAs using Habitat as the only predictor. (b) Results for GEAs using Habitat and the (uninformative) X-coordinate predictor. Results are presented for different sampling strategies (R = random, P = pairs, T = transects), demographies (1R and 2R = refugial expansion, IBD = equilibrium isolation by distance, IM = equilibrium island model), and sample sizes (rows).

### Correction for population structure in RDA

No MEM-based corrections for RDA were applied to IM scenarios, due to low spatial structure (i.e., no PCoA axes were retained based on the broken-stick criterion). The more liberal approach to correction using MEMs (removing retained MEMs significantly correlated with environment), resulted in removal of MEMs with correlation coefficients ranging from 0.07 to 0.72. Ancestry-based corrections were only applied to IM scenarios with 20 individuals since 6 individual samples had K=1. All approaches that correct for population structure in RDA resulted in substantial loss of power across all scenarios, both in terms of empirical *p*-values and cutoff TPRs (Table 1 and Table S4). False positive rates (which were already very low for RDA) increased slightly when correcting for population structure. There were only two scenarios where FPRs improved (one and two fewer FP detections); however, these scenarios saw a reduction in TPR of 81% and 92%, respectively (Table S4).

**Table 1.** Average change in power (from empirical *p*-values) and true and false positive rates (from cutoffs) for RDA using three different approaches for partialling out population structure. All approaches led to an overall loss of power and an increase in false positive rates. There are no MEM corrections for the IM demography, which has no significant spatial structure. Ancestry corrections apply only to 20 individual runs, where K *≠*1.

| Indiv./<br>deme | Demo-<br>graphy | Ancestry | MEMs uncorr.<br>Habitat | All retained<br>MEMs |
| --- | --- | --- | --- | --- |
| | | Change in power (empirical $p$-values) | | |
| 6 | 1R | -0.53 | -0.59 | -0.72 |
|  | 2R | -0.81 | -0.53 | -0.84 |
|  | IBD | -0.94 | -0.75 | -0.96 |
|  | IM | - | - | - |
| 20 | 1R | -0.26 | -0.14 | -0.58 |
|  | 2R | -0.64 | -0.12 | -0.70 |
|  | IBD | -0.93 | -0.69 | -0.93 |
|  | IM | -0.70 | - | - |
| Average |  | -0.69 | -0.47 | -0.79 |
| Change in TPR (cutoffs) |  |  |  |  |
| 6 | 1R | -0.39 | -0.43 | -0.69 |
|  | 2R | -0.70 | -0.40 | -0.76 |
|  | IBD | -0.93 | -0.69 | -0.94 |
|  | IM | - | - | - |
| 20 | 1R | -0.16 | -0.16 | -0.47 |
|  | 2R | -0.47 | -0.17 | -0.51 |
|  | IBD | -0.92 | -0.60 | -0.90 |
|  | IM | -0.71 | - | - |
| Average |  | -0.61 | -0.41 | -0.71 |
| Change in FPR (cutoffs) |  |  |  |  |
| 6 | 1R | 0.0011 | 0.0013 | 0.0020 |
|  | 2R | 0.0021 | 0.0011 | 0.0021 |
|  | IBD | 0.0025 | 0.0017 | 0.0023 |
|  | IM | - | - | - |
| 20 | 1R | 0.0005 | 0.0003 | 0.0014 |
|  | 2R | 0.0014 | 0.0003 | 0.0015 |
|  | IBD | 0.0021 | 0.0010 | 0.0021 |
|  | IM | 0.0023 | - | - |
| Average |  | 0.0017 | 0.0009 | 0.0019 |

**Figure 8.**
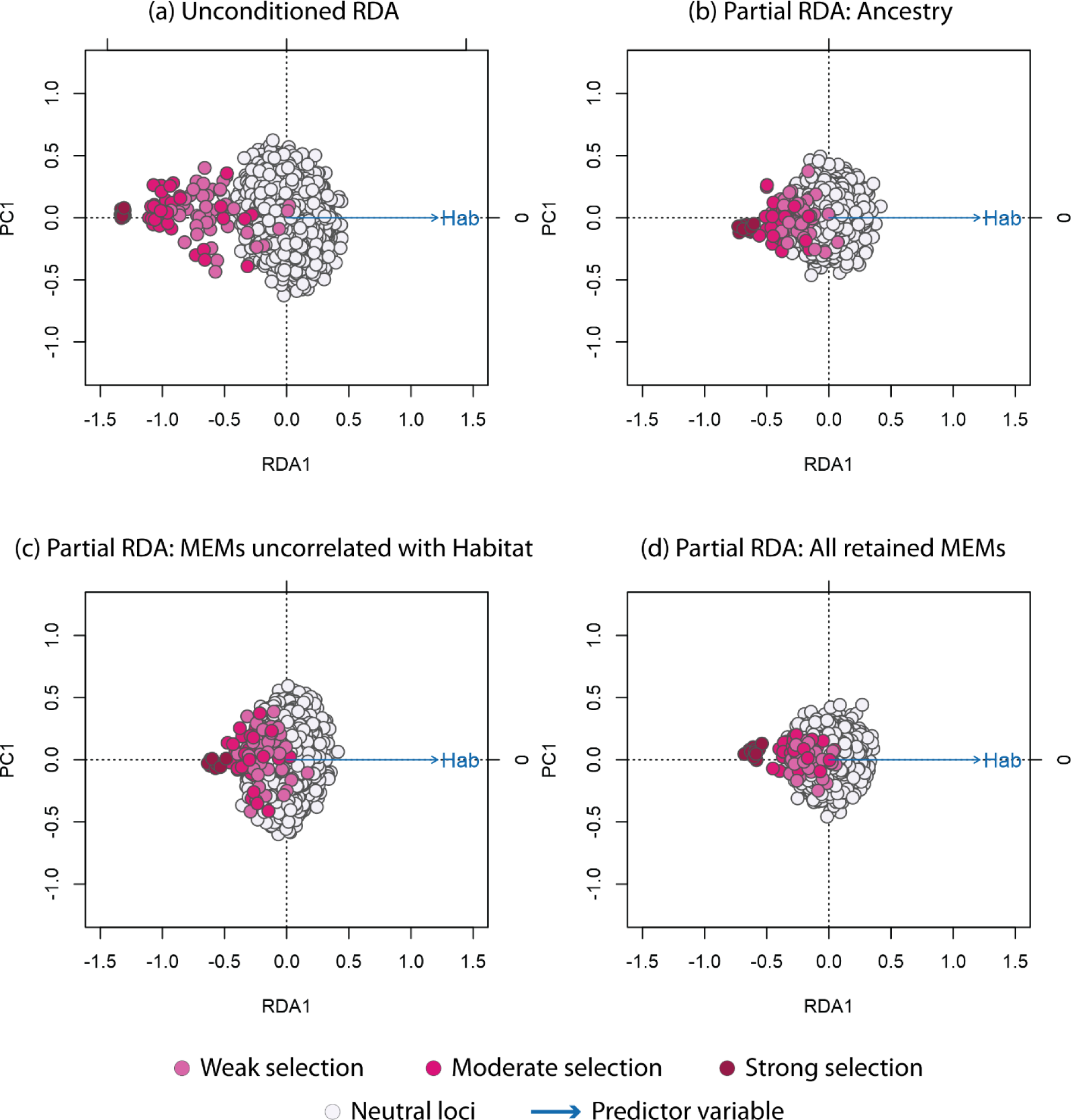
Redundancy analysis biplots for simulation 1R, paired sampling, environmental surface 453, and 6 individuals per deme. Distribution of loci using: (a) unconditioned RDA (no correction for population structure); (b) partial RDA using ancestry values; (c) partial RDA using retained MEMs that are not significantly correlated with Habitat; (d) partial RDA using all retained MEMs.

### Empirical data set

There were four significant RDA axes in the ordination of the wolf data set (Fig. 9), which returned 556 unique candidate loci that loaded +/− 3 SD from the mean loading on each axis: 171 SNPs detected on RDA axis 1, 222 on RDA axis 2, and 163 on RDA axis 3 (Fig. 10). Detections on axis 4 were all redundant with loci already identified on axes 1-3. The majority of detected SNPs were most strongly correlated with precipitation covariates: 231 SNPs correlated with annual precipitation (AP) and 144 SNPs correlated with precipitation seasonality (cvP). The number of SNPs correlated with the remaining predictors were: 72 with mean diurnal temperature range (MDR); 79 with annual mean temperature (AMT); 13 with NDVI; 12 with elevation; 4 with temperature seasonality (sdT); and 1 with percent tree cover (Tree).

**Figure 9.**
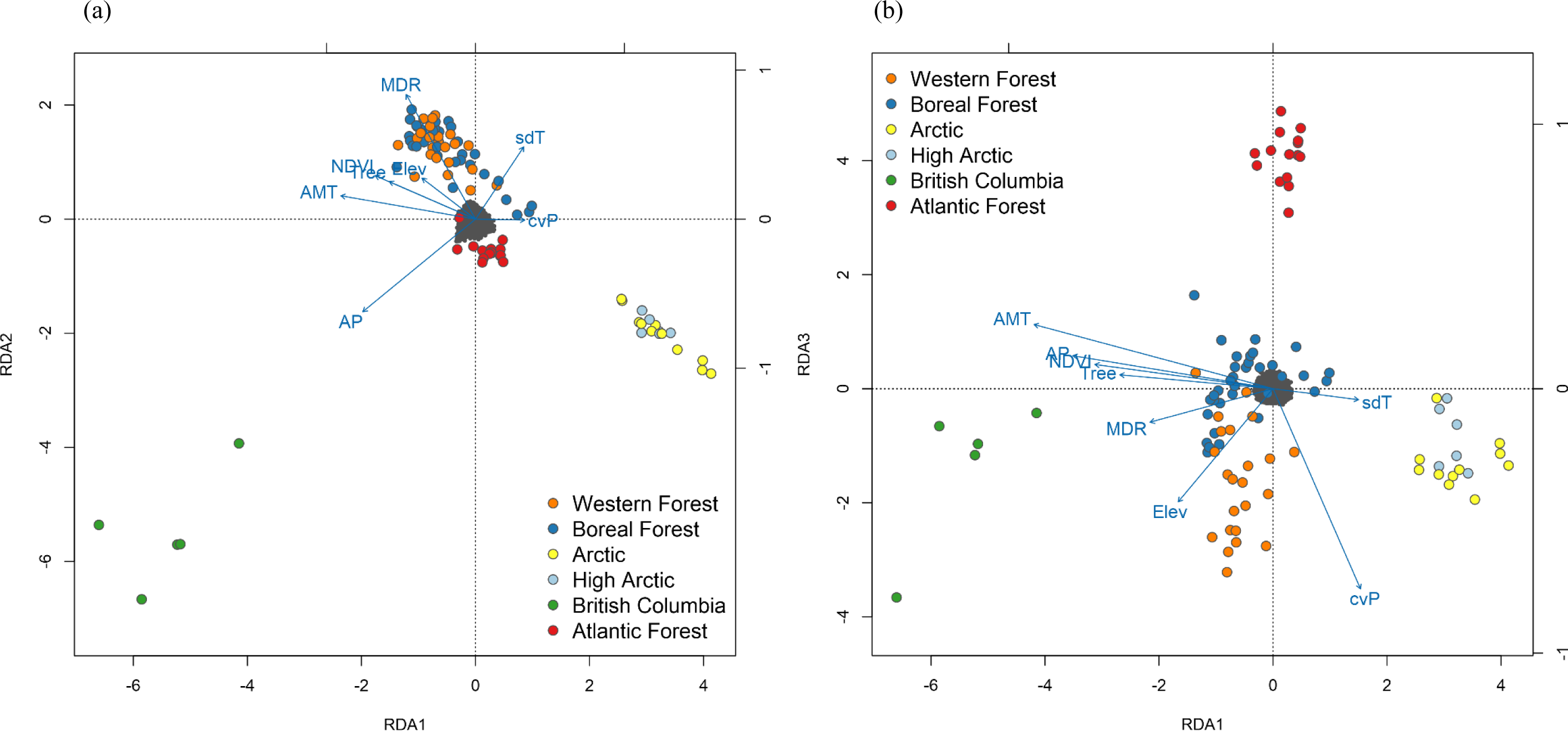
Triplots of wolf data for (a) RDA axes 1 and 2, and (b) axes 1 and 3. The dark gray cloud of points at the center of each plot represent the SNPs, colored points represent individual wolves with coding by ecotype. Blue vectors represent environmental predictors (see text for abbreviations). Triplot scaling is symmetrical (both SNP and individual scores are scaled symmetrically by the square root of the eigenvalues).

**Figure 10.**
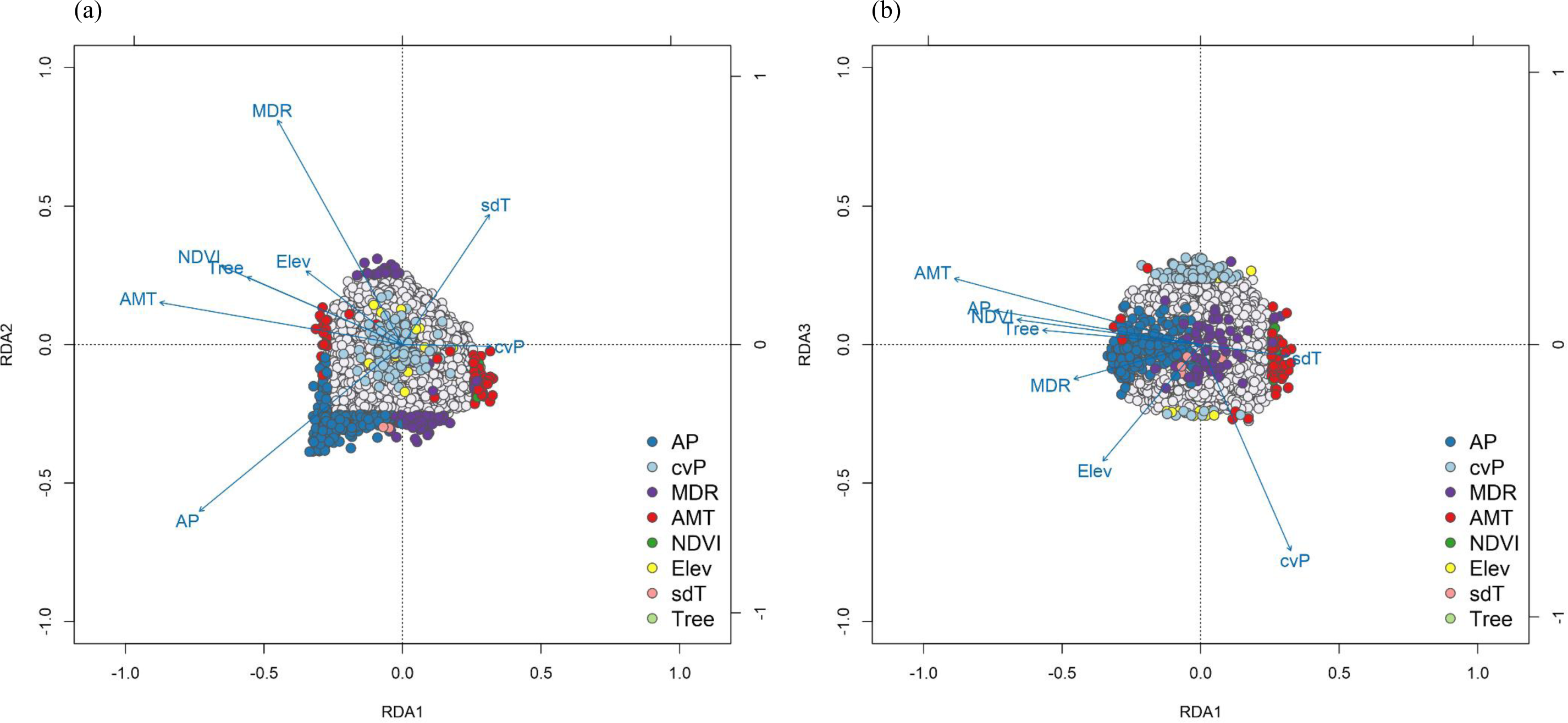
Magnification of wolf data triplots from Figure 9 to highlight SNP loadings on (a) RDA axes 1 and 2, and (b) axes 1 and 3. Candidate SNPs are shown as colored points with coding by most highly correlated environmental predictor. SNPs not identified as candidates (neutral SNPs) are shown in light gray. Blue vectors represent environmental predictors (see text for abbreviations).

## Discussion

Multivariate genotype-environment association (GEA) methods have been noted for their ability to detect multilocus selection (Rellstab *et al.,* 2015; Hoban *et al.,* 2016), although there has been no controlled assessment of the effectiveness of these methods in detecting multilocus selection to date. Since these approaches are increasingly being used in empirical analyses (e.g. Bourret et al. 2014; Brieuc et al. 2015; Pavey et al. 2015; Hecht et al. 2015; Laporte et al. 2016; Brauer et al. 2016), it is important that these claims are evaluated to ensure that the most effective GEA methods are being used, and that their results are being appropriately interpreted.

Here we compare a suite of methods for detecting selection in a simulation framework to assess their ability to correctly detect multilocus selection under different demographic and sampling scenarios. We found that constrained ordinations had the best overall performance across the demographies, sampling designs, sample sizes, and selection levels tested here. The univariate LFMM method also performed well, though power was scenario-dependent and was reduced for loci under weak selection (in agreement with findings by de Villemereuil *et al.,* 2014). Random Forest, by contrast, had lower detection rates overall. In the following sections we discuss the performance of these methods and provide suggestions for their use on empirical data sets.

## Random Forest

Random Forest performed relatively poorly as a GEA. This poor performance is caused by the sparsity of the genotype matrix (i.e., most SNPs are not under selection), which results in detection that is dominated by strongly selected loci (i.e., loci with strong marginal effects). This issue has been documented in other simulation and empirical studies (Goldstein *et al.,* 2010; Winham *et al.,* 2012; Wright *et al.,* 2016) and indicates that RF is not suited to identifying weak multilocus selection or interaction effects in these large data sets. Empirical studies that have used RF as a GEA have likely identified a subset of loci under strong selection, but are unlikely to have identified loci underlying more complex genetic architectures. Note that the amount of environmental variance explained by the RF model can be high (i.e., overall percent variance explained by the detected SNPs, which ranged from 79-91% for these simulations, Table S5), while still failing to identify most of the loci under selection. Removing strong associations from the genotypic matrix can potentially help with the detection of weaker effects (Goldstein *et al.,* 2010), but this approach has not been tested on large matrices. Combined with the computational burden of this method (taking ∼10 days on a single core for the larger data sets), as well as the availability of fast and accurate alternatives such as RDA (which takes ∼3 minutes on the same data), it is clear that RF is not a viable option for GEA analysis of genomic data.

Random Forest does hold promise for the detection of interaction effects in much smaller data sets (e.g., tens of loci, Holliday *et al.* 2012). However, this is an area of active research, and the capacity of RF models in their current form to both capture and identify SNP interactions has been disputed (Winham *et al.,* 2012; Wright *et al.,* 2016). New modifications of RF models are being developed to more effectively identify interaction effects (e.g. Li et al. 2016), but these models are computationally demanding and are not designed for large data sets. Overall, extensions of RF show potential for identifying more complex genetic architectures on small sets of loci, but caution is warranted in using them on empirical data prior to rigorous testing on realistic simulation scenarios.

## Constrained ordinations

The three constrained ordination methods all performed well. RDA in particular had the highest overall power across all methods tested here (Figs. 1-3). Ordinations were relatively insensitive to sample size (6 vs 20 individuals sampled per deme), with the exception of the IM demography, where larger sample sizes consistently improved TPRs, as previously noted by De Mita *et al.* (2013) and Lotterhos & Whitlock (2015) for univariate GEAs. Power was lowest in the IM demography, which is typified by a lack of spatial autocorrelation in allele frequencies and a reduced signal of local adaptation (Table S6), making detection more difficult. This corresponds with univariate GEA results from Lotterhos & Whitlock (2015), who found very low detection rates for loci under weak selection in the IM demography. Power was highest for IBD, followed by the 2R and 1R demographies. Data from natural systems likely lie somewhere among these demographic extremes, and successful differentiation in the presence of IBD and non-equilibrium conditions indicate that ordinations should work well across a range of natural systems.

All three methods were relatively insensitive to sampling design, with transects performing slightly better in 1R and random sampling performing worst in IM (Figs. 4, 6, and S2). Otherwise results were consistent across designs, in contrast to the univariate GEAs tested by Lotterhos and Whitlock (2015), most of which had higher power with the paired sampling strategy. Ordinations are likely less sensitive to sampling design since they take advantage of covarying signals of selection across loci, making them more robust to sampling that does not maximize environmental differentiation (e.g., random or transect designs). All methods performed similarly in terms of detection rates across selection strengths (Figs. 4 and S2). As expected, weak selection was more difficult to detect than moderate or strong selection, except for IBD, where detection levels were high regardless of selection.

High TPRs were maintained when using cutoffs for all three ordination methods (Fig. 6). False positives were universally low for RDA and dbRDA. By contrast, cRDA showed high FPRs for all demographies except IM, tempering its slightly higher TPRs. These higher FPRs are a consequence of using component axes as predictors. Across all scenarios and sample sizes, cRDA detected component 1, 2, or both as significantly associated with the constrained RDA axes (Table S7). Most selected loci load on these components (keeping TPRs high), but neutral markers also load on these axes, especially in cases where there are strong trends in neutral loci (i.e., maximum trends in neutral markers reflect FPRs for cRDA, Table S6, Fig. 6). Given these results, we hypothesized that it might be challenging for cRDA to detect weak selection in the absence of a covarying signal from loci with stronger selection coefficients. If the selection signature is weak, it may load on a lower-level component axis (i.e., an axis that explains less of the genetic variance), or it may load on higher-level axes, but fail to be significantly associated with the constrained axes. Note that although cRDA contains a step to reduce the number of components, parallel analysis resulted in retention of all axes in every simulation tested here (Table S7). This meant that cRDA could search for the signal of selection across all possible components.

When tested on simulations with loci under weak selection only, RDA, which uses the genotype matrix directly, maintained similar power as in the full data set (except in the IM scenario, where power was higher when all selected loci were included), indicating that selection signals can be detected with this method in the absence of loci under strong selection (Fig. 5, top row). By contrast, cRDA detection was more variable, ranging from comparable detection rates with the full data set, to no/poor detections under certain demographies and sample sizes. In these latter cases, poor performance is reflected in the component axes detected as significant (Table S7); instead of identifying the signal in the first few axes, a variable set of lower-variance axes are detected (or none are detected at all). This indicates that the method is not able to “find” the selected signal in the component axes in cases where that signal is not driven by strong selection. This result, in addition to higher FPRs for cRDA, builds a case for using the genotype matrix directly with a constrained ordination such as RDA or dbRDA, as opposed to a preliminary step of data conversion with PCA.

## Should results from different tests be combined?

A common approach in local adaptation studies is to run multiple tests (GEA only, or a combination of GEA and differentiation methods) and look for overlapping detections across methods. This ad hoc approach is thought to increase confidence in TPRs, while minimizing FPRs. The problem with this approach is that it can bias detection toward strong selective sweeps to the exclusion of other adaptive mechanisms which may be equally important in shaping phenotypic variation (Le Corre & Kremer, 2012; François *et al.,* 2016). If the goal is to detect other forms of selection such as recent selection or selection on standing genetic variation, this approach will not be effective since most methods are unlikely to detect these weak signals. Additionally, this approach limits detections to those of the least powerful method used, forcing overall detection rates to be a function of the weakest method implemented.

The complexities of this issue are illustrated by comparing results across two sets of RDA and LFMM results: one where the driving environmental variable is known (Fig. 7a), and another where the environmental predictors represent hypotheses about the most important factors driving selection (Fig. 7b). In both cases, agreement on TPs is high, and RDA has a large number of true positive detections that are unique to that method, while unique detections by LFMM are largely limited to the IM demography. The differences in the cases lies in FP detections: when selection is well understood, and uninformative predictors are not used, retaining RDA detections only is the approach that will maximize TPRs (and detection of weak loci under selection) while maintaining minimal to zero FPRs (Fig. 7a). Where GEA analyses are more exploratory (i.e., when selective gradients are unknown), combining detections can help reduce FPRs (Fig. 7b). If some FP detections are acceptable, keeping only RDA detections will improve TPRs at the cost of slightly increased FPRs. A third approach, keeping all detections across both methods, would yield little improvement in TPRs in both cases, since LFMM has high levels of unique FPs, and minimal unique TP detections.

The decision of whether and how to combine results from different tests will be specific to the study questions, the tolerance for false negative and false positive detections, and the capacity for follow-up analyses on detected markers. For example, if the goal is to detect loci with strong effects while keeping false positive rates as low as possible, or GEA is being used as an exploratory analysis, running multiple GEA methods and considering only overlapping detections could be a suitable strategy. However, if the goal is to detect selection on standing genetic variation or a recent selection event, and the most important selective agents (or close correlates of them) are known, combining detections from multiple tests would likely be too conservative. In this case, the best approach would be to use a single GEA method, such as RDA, that can effectively detect covarying signals arising from multilocus selection, while being robust to selection strength, sampling design, and sample size.

## Correction for population structure

All three methods used to correct for populations structure in RDA resulted in substantial loss of power and, in most cases, increased FPRs (Table 1 and S4). The effect of correcting for population structure can be seen in ordination biplots from an example simulation scenario (Fig. 8). In this 1R demographic scenario, the selection surface (“Hab”) and the refugial expansion gradient coincide, so any correction for population structure will also remove the signal of selection from the selected loci. The correction is most conservative when using all significant MEM predictors to account for spatial structure (Fig. 8d), and is less conservative when using only MEMs not significantly correlated with environment (Fig. 8c), or ancestry coefficients (Fig. 8b). In all cases, however, the loss of the selection signal is significant (Table 1), and is visible in the increasing overlap of selected loci with neutral loci.

While the simulations used here have overall low global Fst (average Fst = 0.05), population structure is significant enough in many scenarios to result in elevated FPRs for GLMs (univariate linear models which do not correct for population structure, Fig. 6). Despite this, RDA and dbRDA (the multivariate analogue of GLMs) do not show elevated FPRs, even when selection covaries with a range expansion front, as in the 1R and 2R demographies. This is likely because only loci with extreme loadings are identified as potentially under selection, leaving most neutral loci, which share a similar, but weaker, spatial signature, loading less than +/− 3 SD from the mean. The generality of these results needs to be tested in a comprehensive manner using an expanded simulation parameter space that includes stronger population structure and metapopulation dynamics; this work is currently in progress. In the meantime, we recommend that RDA be used conservatively in empirical systems with higher population structure than is tested here, for example, by finding overlap between detections identified by RDA and LFMM (or another GEA that accounts for population structure).

## Empirical example

Triplots of three of the four significant RDA axes for the wolf data show SNPs (dark gray points), individuals (colored circles), and environmental variables (blue arrows, Fig. 9). The relative arrangement of these items in the ordination space reflects their relationship with the ordination axes, which are linear combinations of the predictor variables. For example, individuals from wet and temperate British Columbia are positively related to high annual precipitation (AP) and low temperature seasonality (sdT, Fig. 9a). By contrast, Artic and High Arctic individuals are characterized by small mean diurnal temperature range (MDR), low annual mean temperature (AMT), lower levels of tree cover (Tree) and NDVI (a measure of vegetation greenness), and are found at lower elevation (Fig. 9a). Atlantic Forest and Western Forest individuals load more strongly on RDA axis 3, showing weak and strong precipitation seasonality (cvP) respectively (Fig. 9b), consistent with continental-scale climate in these regions.

If we zoom into the SNPs, we can visualize how candidate SNPs load on the RDA axes (Fig. 10). For example, SNPs most strongly correlated with AP have strong loadings in the lower left quadrant between RDA axes 1 and 2 along the AP vector, accounting for the majority of these 231 AP-correlated detections (Fig. 10a). Most candidates highly correlated with AMT and MDR load strongly on axes 1 and 2, respectively. Note how candidate SNPs correlated with precipitation seasonality (cvP) and elevation are located in the center of the plot, and will not be detected as outliers on axes 1 or 2 (Fig. 10a). However, these loci are detected as outliers on axis3 (Fig. 10b). Overall, candidate SNPs on axis 1 represent multilocus haplotypes associated with annual precipitation and mean diurnal range; SNPs on axis 2 represent haplotypes associated with annual precipitation and annual mean temperature; and SNPs on axis 3 represent haplotypes associated with precipitation seasonality.

Of the 1661 candidate SNPs identified by Schweizer *et al.,* (2016) using Bayenv (Bayes Factor > 3), only 52 were found in common with the 556 candidates from RDA. Of these 52 common detections, only nine were identified based on the same environmental predictor. If we include Bayenv detections using highly correlated predictors (removed for RDA) we find nine more candidates identified in common. Additionally, only 18% of the Bayenv identifications were most strongly related to precipitation variables, which are known drivers of morphology and population structure in gray wolves (Geffen *et al.,* 2004; O’Keefe *et al.,* 2013; Schweizer *et al.,* 2016). By contrast, 67% of RDA detections were most strongly associated with precipitation variables, providing new candidate regions for understanding local adaptation of gray wolves across their North American range.

## Conclusions and recommendations

We found that constrained ordinations, especially RDA, show a superior combination of low FPRs and high TPRs across weak, moderate, and strong multilocus selection. These results were robust across the levels of population structure, demographic histories, sampling designs, and sample sizes tested here. Additionally, RDA outperformed an alternative ordination-based approach, cRDA, especially (and importantly) when the multilocus selection signature was completely derived from loci under weak selection. It is important to note that population structure was relatively low in these simulations. Results may differ for systems with strong population structure or metapopulation dynamics, where it can be important to correct for structure or combine detections with another GEA that accounts for structure. Continued testing of these promising methods is needed in simulation frameworks that include more population structure, multiple selection surfaces, and genetic architectures that are more complex than the multilocus selection response modeled here. However, this study indicates that constrained ordinations are an effective means of detecting adaptive processes that result in weak, multilocus molecular signatures, providing a powerful tool for investigating the genetic basis of local adaptation and informing management actions to conserve the evolutionary potential of species of agricultural, forestry, fisheries, and conservation concern.

## Acknowledgements

We thank Katie Lotterhos for sharing her simulation data (Lotterhos & Whitlock 2015) and for additional spatial coordinate data and code. Tom Milledge with Duke Resource Computing provided invaluable assistance with the Duke Compute Cluster. We also thank Olivier François for helpful advice with LFMM, and three reviewers for constructive feedback that greatly improved the manuscript. BRF was supported by a Katherine Goodman Stern Fellowship from the Duke University Graduate School and a PEO Scholar Award.

## Data accessibility

Simulation data from Lotterhos & Whitlock (2015): Dryad: doi:10.5061/dryad.mh67v Supporting simulation data (coordinate files) for Lotterhos & Whitlock (2015) data provided by Wagner *et al.* (2017): Dryad: doi:10.5061/dryad.b12kk. Wolf data from Schweizer *et al.* (2016): Dryad: doi.org/10.5061/ dryad.c9b25.

## Author contributions

BRF and DLU conceived the study. BRF performed the analyses and wrote the manuscript. HHW contributed code. JRL, HHW, and DLU helped interpret the results and write the manuscript.

## Supporting information

Forester_Simulation_Rcode.zip: Contains R code for data preparation and all of the methods tested.

Forester_Wolf_Rcode.zip: Contains R code for data preparation, analysis, interpretation, and plotting of the wolf data set with RDA.

Forester_SI.pdf: Supplemental Figures S1-S7 and Tables S1-S7.

